# Dietary folic acid deficiency impacts hippocampal morphology and cortical acetylcholine metabolism in adult male and female mice

**DOI:** 10.1101/2020.12.13.422535

**Authors:** Calli Bennett, Jacalyn Green, Mae Ciancio, Joanna Goral, Lenore Pitstick, Matthew Pytynia, Alice Meyer, Neha Kwatra, Nafisa M. Jadavji

**Affiliations:** Biomedical Sciences Program, College of Graduate Studies, Midwestern University, Glendale, US; College of Osteopathic Medicine, Midwestern University, Glendale, US; Biochemistry and Molecular Genetics, College of Graduate Studies, Midwestern University, Downers Grove, US; Biomedical Sciences Program, College of Graduate Studies, Midwestern University, Downers Grove, US; Anatomy, College of Graduate Studies, Midwestern University, Downers Grove, US; College of Dental Medicine, Midwestern University, Glendale, US; College of Veterinary Medicine, Midwestern University, Glendale, US; Department of Neuroscience, Carleton University, Ottawa, Canada

**Keywords:** One- carbon metabolism, folic acid, hippocampus, apoptosis, acetylcholine, sex differences

## Abstract

**Objective:** One-carbon metabolism is a metabolic network that integrates nutritional signals with biosynthesis, redox homeostasis, and epigenetics. There are sex differences in hepatic one-carbon metabolism. However, it is unclear whether there are sex differences in dietary deficiencies of one-carbon metabolism in the brain. The aim of this study was to investigate if sex modulates the effects of dietary folic acid deficiency in brain tissue using a mouse model.

**Methods:** Male and female C57Bl/6J mice were placed on a folic acid deficient (FD) or control diet (CD) at six weeks of age. Mice were maintained on these diets for six months, after which animals were euthanized and brain tissue and serum were collected for analysis. Serum folate levels were measured. In brain tissue, hippocampal volume and morphology including Cornu Ammonis 1 and 3 (CA1; CA3), and dentate gyrus thickness were measured. Apoptosis within the hippocampus was assessed using active caspase-3 immunofluorescence staining. Additionally, cortical acetylcholine metabolism was measured in brain tissue using immunofluorescence staining of acetylcholinesterase (AChE), or choline acetyltransferase (ChAT), and neuronal nuclei (NeuN).

**Results:** Male and female FD mice had reduced serum levels of folate. Both males and females maintained on a FD showed a decrease in the thickness of the hippocampal CA1-CA3 region. Interestingly, there was a sex difference in the levels of active caspase-3 within the CA3 region of the hippocampus. In cortical tissue, there were increased levels of neuronal ChAT and reduced levels of AChE in FD females and male mice.

**Conclusions:** The results indicated that FD impacts hippocampal morphology and cortical neuronal acetylcholine metabolism. The data from our study indicate that there was only one sex difference and that was in hippocampal apoptosis. Our study provides little evidence that sex modulates the effects of dietary folate deficiency on hippocampal morphology and cortical neuronal acetylcholine metabolism.

## 1. Introduction

One-carbon (1C) metabolism is a key metabolic network that integrates nutrition signals with biosynthesis, redox homeostasis, and epigenetics. It plays an essential role in the regulation of cell proliferation, stress resistance, and embryonic development. Folate, also known as vitamin B9, is the main component of 1C metabolism. Folates act as 1C donors in the regeneration of methionine from homocysteine, nucleotide synthesis and repair, and DNA methylation (1).

While gastrointestinal bacteria produce small amounts of folate, the majority are obtained in our diets. However, the bioavailability and stability from these dietary sources are much lower than synthetic, folic acid (2). Many countries fortify foods with folic acid in order to reduce deficiencies and related negative health outcomes (3). However, even in countries in which fortifying foods is routine, deficiencies remain fairly common. Deficiencies are seen in individuals that have a high rate of tissue turnover (e.g., pregnant women), have an alcohol use disorder (4), or in those using folate antagonist drugs (e.g., methotrexate) (5). Additionally, deficiencies are common in the elderly population, which is especially vulnerable to malnutrition and folate absorption impairments (6).

Folate’s role in the regeneration of methionine from homocysteine proves especially significant when considering vascular and brain health. Elevated levels of homocysteine, that can be a result of folic acid deficiency, have been linked to several negative health outcomes, including increased risk for Alzheimer’s Disease (7), and cerebrovascular diseases including stroke (8). Additionally, hyperhomocysteinemia is an important indicator of post-stroke recovery (9), mortality (10), and risk of recurrence (11). Homocysteine leads to downstream oxidative damage (12), indicating a potential mechanism behind its role in neuronal degeneration and atherosclerosis (13).

Choline, another component of 1C metabolism, is an essential nutrient. Folate and choline metabolism are tightly linked. In model systems that study the impact of reduced levels of dietary folates, choline metabolism is also affected (14–17). Choline can act as a 1C donor, especially in situations of diminished folate. In the brain, choline is also involved in the production of acetylcholine, the main neurotransmitter of the parasympathetic nervous system, and lipid membrane synthesis. Acetylcholine is synthesized by choline acetyltransferase (ChAT) and hydrolyzed by acetylcholinesterase (AChE). Reduced levels of choline have been reported to result in more apoptosis in cultured cells (18) and were observed in the hippocampus of offspring of choline deficient females (19). *In vivo* rodent studies have also shown cognitive impairment, because of a dietary choline deficiency. The hippocampus is particularly vulnerable to both choline and folate dietary deficiencies (14,16,20,21).

There is growing evidence that there are significant sex differences with regard to 1C metabolism (22–24), including quantifiable differences in homocysteine levels (25). Males homozygotic for the polymorphism in methylenetetrahydrofolate reductase are more vulnerable to factors (e.g., smoking) that increase levels of homocysteine (25). Estrogen may serve as a protective factor because it is linked to reduced homocysteine levels in blood. Decreased homocysteine levels lead to declined risk for vascular diseases (25). Furthermore, using a model, five key hepatic 1C metabolism enzymes are differentially expressed in males and females(23). Estrogen and other female hormones impact expression of these enzymes, leading to increased levels of homocysteine in males, and increased levels of choline and betaine in females(23). Further, folate intake and expression of genes in 1C metabolism regulate steatosis in a sex specific manner (26). Despite the significant evidence of hepatic 1C differences between sexes, limited evidence exists on the brain. A recent study, including both male and female mice, demonstrated that there are significant sex differences in behavior observed caused by a methylenetetrahydrofolate reductase deficiency (22,24). The aim of this study was to investigate if sex modulates the effects of dietary folic acid deficiency in brain tissue using a mouse model. We hypothesized dietary folic acid deficiency would result in reduced levels dietary folate in males and females and that sex would impact on hippocampal morphology and cortical acetylcholine metabolism in mice maintained on a folic acid deficient diet.

## 2. Methods

### 2.1 Experimental design

All experiments were done in accordance with IACUC animal welfare protocols and were approved by the Midwestern University Downers Grove IACUC Committee. Male and female C57Bl/6J mice were put on a folic acid deficient (FD) or control (CD) diets (Envigo) at 6 weeks of age (Table 1). Animals had access to food and water *ad libitum*. Each diet group had 5 males and 5 females; a total of 20 animals were used.

**Table 1.** List of micro- and macro-nutrient contents in Envigo control (TD.04194) and folic acid deficient diet (TD.95247). The Envigo folic acid deficient (TD.95247) was identical to the control diet (TD.04194) except it contained 0.2 mg/kg folic acid.

| Category | Concentration | Category | Concentration |
| --- | --- | --- | --- |
| **Amino acids** | **(g/kg)** | **Lipids** | **(g/kg)** |
| Lysine | 14 | Total fat | 60.0 |
| Methionine | 4.7 | Saturated fat | 9.0 |
| Cystine | 3.6 | Monounsaturated fat | 13.8 |
| Arginine | 6.6 | Polyunsaturated fat | 36.6 |
| Phenylalanine | 8.8 | 4:0 Butyric acid | 0.0 |
| Tyrosine | 9.2 | 6:0 Caproic Acid | 0.0 |
| Histidine | 5.1 | 8:0 Caprylic Acid | 0.0 |
| Isoleucine | 10.1 | 10:0 Capric acid | 0.0 |
| Leucine | 16 | 12:0 Lauric acid | 0.0 |
| Threonine | 7.6 | 14:0 Myristic Acid | 0.0 |
| Tryptophan | 2.1 | 16:0 Palmitic acid | 6.6 |
| Valine | 11.7 | 16:1 Palmitoleic acid | 0.0 |
| Aspartic Acid | 12.1 | 18:0 Stearic acid | 2.4 |
| Glutamic acid | 36.6 | 18:1 Oleic acid | 13.8 |
| Alanine | 5.2 | 18:2 Linoleic acid | 31.8 |
| Glycine | 3.2 | 18:3 Linolenic acid | 4.8 |
| Proline | 18.1 | Cholesterol | 8.8 mg/kg |
| Serine | 10 |  |  |
| **Fat-soluble Vitamins** | **(IU/kg)** | **Minerals** | **(mg/kg)** |
| Vitamin A | 4000 | Calcium | 5100 |
| Vitamin D | 1000 | Phosphorus | 3000 |
| Vitamin E | 75 | Potassium | 3600 |
| Vitamin K | 0.8 | Sodium | 1000 |
| **Water-soluble Vitamins** | **(mg/kg)** | Chlorine | 1600 |
| Biotin | 0.20 | Magnesium | 515 |
| Choline | 1147.5 | Copper | 6.0 |
| Folic acid | 2.0 | Iron | 36.5 |
| Niacin | 30.0 | Zinc | 35.6 |
| Pantothenate | 14.7 | Manganese | 10.5 |
| Riboflavin | 6.0 | Iodine | 0.21 |
| Thiamin | 4.9 | Selenium | 0.15 |
| Vitamin B<sub>6</sub> | 5.8 | Molybdenum | 0.15 |
| Vitamin B<sub>12</sub> | 0.03 | Chromium | 1.00 |
| Vitamin C | 0.0 |  |  |

The CD and FD contained the same concentrations of macro-micro-nutrients, except for folic acid (Table 1). The CD contained 2.0 mg/kg of folic acid (TD.04194), while FD contained 0.2 mg/kg of folic acid (TD.95247) (27,28). Mice were maintained on these diets for 6 months. Animals were weighed weekly throughout the experiment. At 8 months of age mice were euthanized with CO_2_ inhalation and cervical dislocation. This corresponds to mature adults in humans (29).

### 2.2 Tissue collection

At time of euthanization body and brain weights were recorded. Tissue including brain and blood were collected for analysis. Serum was isolated from blood and stored at −80°C until analysis. Brain tissues were fixed in 4% paraformaldehyde overnight and then switched over to a 70% ethanol solution. Tissue was processed and embedded in paraffin in a coronal orientation. Brain tissue was sectioned using a microtome (Leica) at 5-μm thickness. Sections were slide mounted and stored at room temperature for staining.

### 2.3 Microbiological assay for serum folate

Serum folate was measured using a microbiological assay (30). *Lactobacillus rhamnosus* (ATCC) was grown overnight in Folic Acid Casei Medium (Difco) supplemented with 0.025% sodium ascorbate (A7631, Sigma-Aldrich,) and 1.2 ng/ml of (6S)-5-formyltetrahydrofolate (Schirks Laboratories). An equal volume of 80% sterile glycerol was added to the overnight culture and 1 ml aliquots were frozen and stored at −80°C. The day of the assay, an aliquot of *L. rhamnosus* was thawed and 100ul was added to 50ml of Folic Acid Casei Medium supplemented with 0.025% sodium ascorbate. The approximately 20 mg/liter folic acid stock (Sigma-Aldrich) was 0.22 μm filter sterilized, verified by absorbance spectra at 282 (molar absorptivity coeffícient= 27,000 M^-1^, MW= 441.4), and stored at −20°C. Dilutions of the folic acid stock and test sera were made in freshly prepared 0.22 μm filter sterilized 0.5% sodium ascorbate. Through a series of dilutions, a folic acid standard curve with 100μl/well in duplicate was generated from 0 to 166 pg/ml final concentration. Test sera was diluted with sterile 0.5% sodium ascorbate before the assay. Several dilutions of sera, 1:600-1:2400 final concentration, at 100μl/well in duplicate were tested to ensure that an A600 fell within the standard curve. The inoculated media was added at 200μl/well to the 10 x10 Bioscreen honeycomb plate (Fisher Scientific). The plate was loaded into the Bioscreen C accompanied with EZExperiment 1.26 software (OY Growth Curves Ab Ltd), incubated at 37°C with shaking for 24 hours, and A_600_ read at 24 hours (30).

### 2.4 Brain tissue morphological analysis

Series of brain tissue samples were stained with 1% cresyl violet (Sigma) to assess morphological changes. Images were taken using Nikon Ni-U Compound Light microscope. Analysis of morphological changes focused on the hippocampal formation as previous studies demonstrate that this area is particularly vulnerable to deficiencies in 1C metabolism (14,16,20). Thickness measurements of the granular cell layer were taken using ImageJ (NIH) (31,32) within the dentate gyrus, Cornu Ammonis 1 (CA1) and 3 (CA3) regions of the hippocampus. The total volume of the hippocampus was measured using ImageJ (NIH) (31,32) by tracing the perimeter of the structure. For each measurement a minimum of 3 brain tissue sections per animals were used and each hemisphere was measured. An average of 6 measurements was used per animal.

### 2.5 Immunofluorescence

Antigen retrieval on paraffin embedded brain tissue sections was performed prior to staining as previously described (33). A series of brain tissue sections were stained with the following antibodies: anti-choline acetyltransferase (ChAT) (Millipore, AB144P) at 1:100; anti-acetylcholinesterase (AChE) (Sigma, SAB2500018) at 1:100; or anti-active caspase-3 (Cell Signaling, 9662) at 1:100. All samples were co-stained with anti-neuronal nuclei antibody (NeuN, ab104224) (AbCam) at 1:200 and (4’,6-diamidino-2-phenylindole) DAPI (Fisher Scientific) at 1:10,000.

Imaging of ChAT and AChE was conducted using a Zeiss Apotome microscope equipped with a camera. Two brain tissue sections with three subfields within each cortex were imaged. Imaging of active caspase-3 was performed using an Olympus inverted microscope equipped with a camera. The hippocampus was analyzed using two images of CA1, CA3, or three dentate gyrus for each animal. Only cells demonstrating colocalization of ChAT, AChE, or active caspase-3 along with NeuN and DAPI were considered positive and counted.

### 2.6 Data and statistical analysis

All data analysis was conducted by 2 individuals blinded to experimental groups. Data were analyzed using GraphPad Prism 9.0 and IBM SPSS Statistics 25. Weekly body weights were analyzed using repeated measures ANOVA using IBM SPSS Statistics 25. Two-way analysis of variance (ANOVA) was used to determine interactions and main effects between diet groups (folic acid deficient or control) and sex (male or female). Analyses included brain and body weight, morphological measurements, immunofluorescence of choline metabolism (AChE and ChAT), and active caspase-3. Pairwise post-hoc testing was performed using the Tukey’s multiple comparison test. In all analyses, p<0.05 was considered statistically significant.

## 3. Results

### 3.1 Body and brain weight

Body weights were collected on a weekly basis; a difference between male and females (Figure 1A, F(_1,14_) = 50.80, p<0.001). However, there was no difference by diet (F(_1,14_) = 0.048, p = 0.830). Brain and body weights were collected at time of euthanization and the data was analyzed. There was a sex difference in body weight (Figure 1B, F (_1, 14_) = 48.48, p < 0.001), but there was no effect of diet (F (_1, 14_) = 1.97, p = 0.18). There was no difference in brain weight observed (Figure 1C, Sex: F (_1, 14_) = 2.58, p = 0.13, Diet: F (_1, 14_) = 1.13, p = 0.31). When analyzing brain weight as a percent of body weight, females had a higher percent brain weight than males (Figure 1D, F (_1, 14_) = 73.99, p < 0.0001). Female FD mice had smaller brain weight as percent of body when compared to female CD mice (F (_1, 14_) = 6.217, p = 0.026).

**Figure 1.**
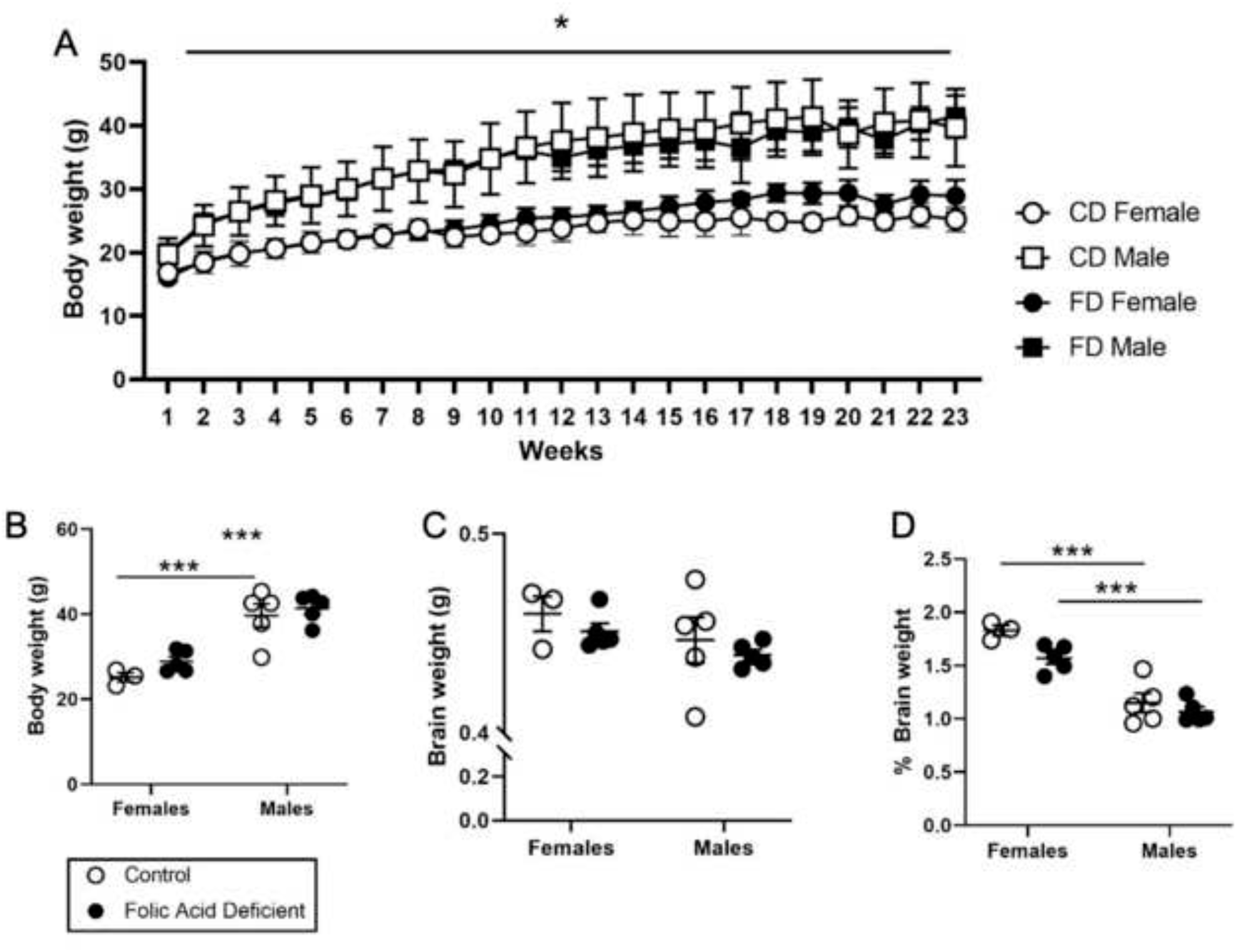
The impact of dietary folic acid deficiency and sex on body and brain weight. Weekly body weights (A) for duration of 6-month experimental period. At time of euthanization body weight (B), brain weight (C) and brain weight as a percent of body weight (D). Line graph and scatter plot with mean + SEM of 3 to 5 mice per group. * p<0.05 indicate sex difference after repeated measures 2-way ANOVA. ***p<0.001 indicate Tukey’s pairwise comparison between groups.

### 3.2 Serum folate measurements

Folate levels were measured in serum collected from all animals to confirm dietary deficiency. There was no sex difference (F (_1,14_) = 2.40, p = 0.14). Compared to CD mice, animals maintained on the FD had reduced levels of folate in their serum (Figure 1A; F (_1,14_) = 10.52, p = 0.0059).

### 3.3 Hippocampal volume and morphology

Previous work has shown that the hippocampus is affected by a 1C deficiency (14,19,21). The hippocampus volume was analyzed using ImageJ, and thickness of the dentate gyrus, CA1, and CA3 regions. There was no sex difference (Figure 2B; F (_1,12_) = 0.37, p = 0.55) in the total hippocampal volume. However, there was a trend for lower hippocampal volume of mice maintained on FD (F (_1,12_) = 4.37, p = 0.059). There was no sex (Figure 2C; F (_1,12_) = 0.86, p = 0.37) or diet (F (_1,12_) = 1.15, p = 0.31) differences in the thickness of the dentate gyrus. There was no sex difference in the thickness of CA1 and 3 regions (F (_1,13_) = 0.94, p = 0.35). However, there was a decrease in the thickness of the CA1 and 3 regions of mice maintained on a FD diet (Figure 2D; F (_1,13_) = 37.3, p < 0.0001).

**Figure 2.**
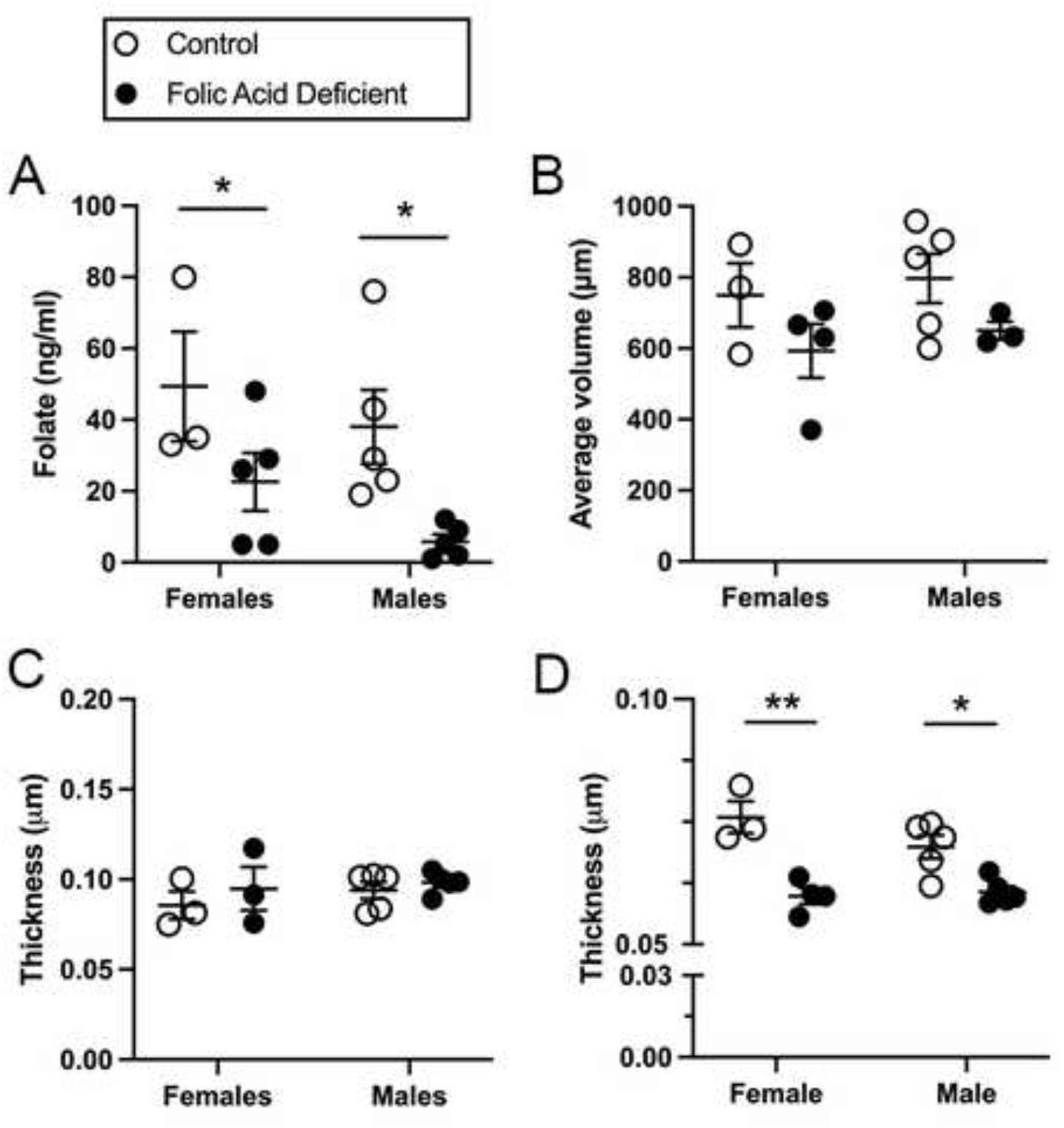
The impact of dietary folic acid deficiency and sex on serum folate levels (A) and hippocampal volume (B), and thickness of dentate gyrus granular cell layer (C) and Cornu Ammonis (CA) 1 and 3 region (D). Scatter plot with mean + SEM of 3 to 5 mice per group. *p<0.05 and **p<0.01 indicate Tukey’s pairwise comparison between groups.

### 3.4 Hippocampal apoptosis

Hippocampal apoptosis was assessed by counting active caspase-3 positive neuronal cells, such cells were counted within the dentate gyrus, CA1, and CA3 regions of the hippocampus. There was no difference in neuronal apoptosis between groups within the dentate gyrus (sex: F (_1,12_) = 0.035, p = 0.86, diet: F (_1,13_) = 0.033, p = 0.86). There was a trend for a sex difference of apoptotic neurons within the CA1 region (F (_1,12_) = 3.502, p = 0.084), but there was no difference between dietary groups (F (_1,12_) = 0.00083, p = 0.99). Representative images of active caspase-3 staining within the CA3 are shown in Figures 3A to D. There was a difference in the number of positive apoptotic neurons between females and males (Figure 3E; (F (_1,12_) = 7.36, p = 0.019)), but no dietary difference (F (_1,12_) = 1.44, p = 0.25).

**Figure 3.**
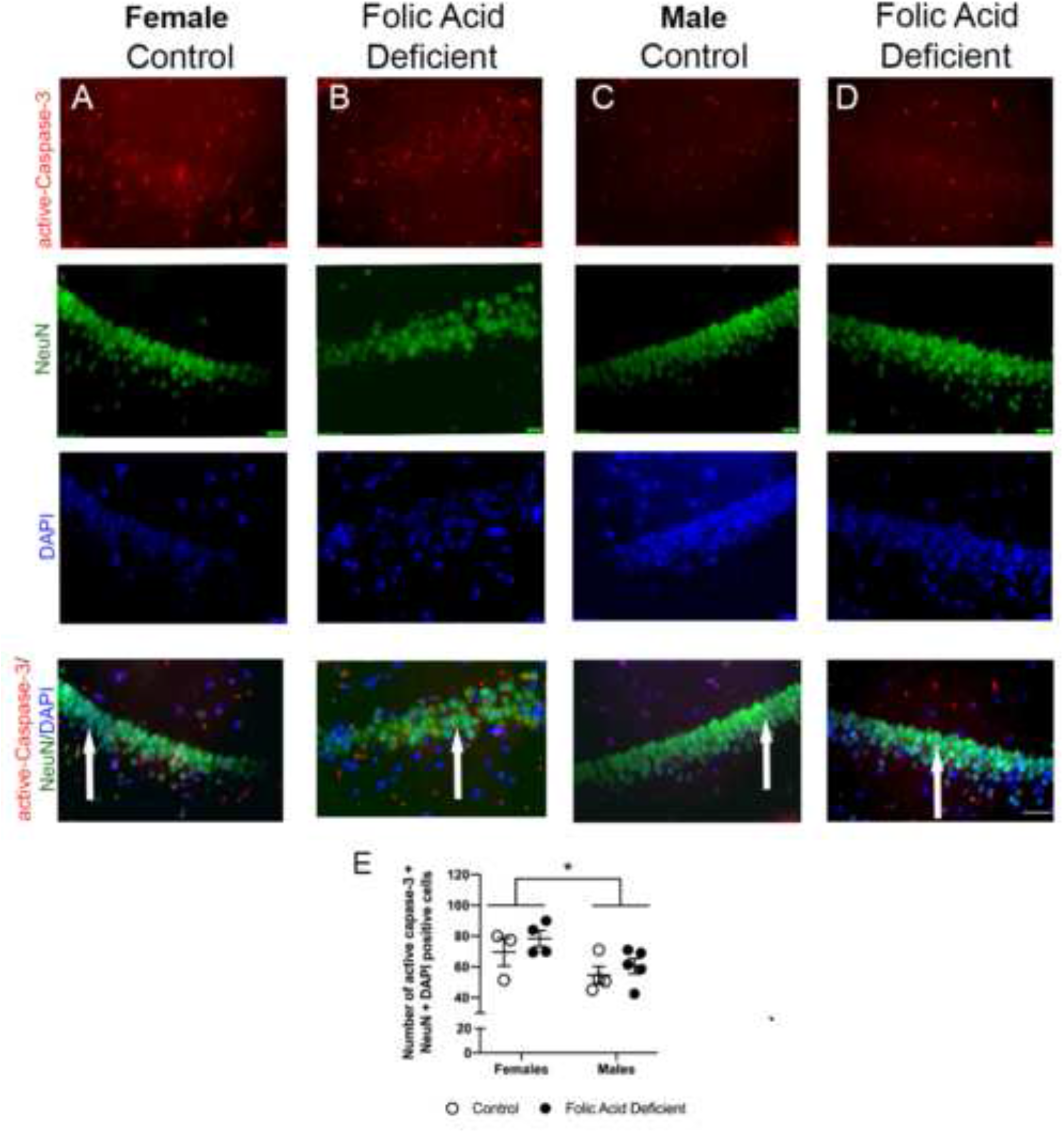
The impact of dietary folic acid deficiency and sex on active caspase-3 levels within the hippocampal Cornu Ammonis 3 (CA3) region of brain tissue. Representative images of active caspase-3, neuronal nuceli (NeuN), and 4’6-diamidino-2-phenyloindole (DAPI) from all experimental groups (A-D). Blinded quantification of active caspase-3, NeuN, and DAPI positive cells (E). Scatter plot with mean + SEM of 3 to 5 mice per group. *p<0.05 indicates sex main effect between males and females.

### 3.5 Cortical choline metabolism

Neuronal cortical choline metabolism was characterized by counting the number choline acetyltransferase (ChAT) positive cells. This enzyme is involved in the synthesis of acetylcholine at the synapse. Representative images are shown in Figure 4A to D. Quantification revealed no difference between sexes (F (_1,12_) = 1.79, p = 0.21), however, there were increases in ChAT positive neurons in the FD groups when compared to CD mice (Figure 4E; F(_1,12_) = 4.58, p = 0.050).

**Figure 4.**
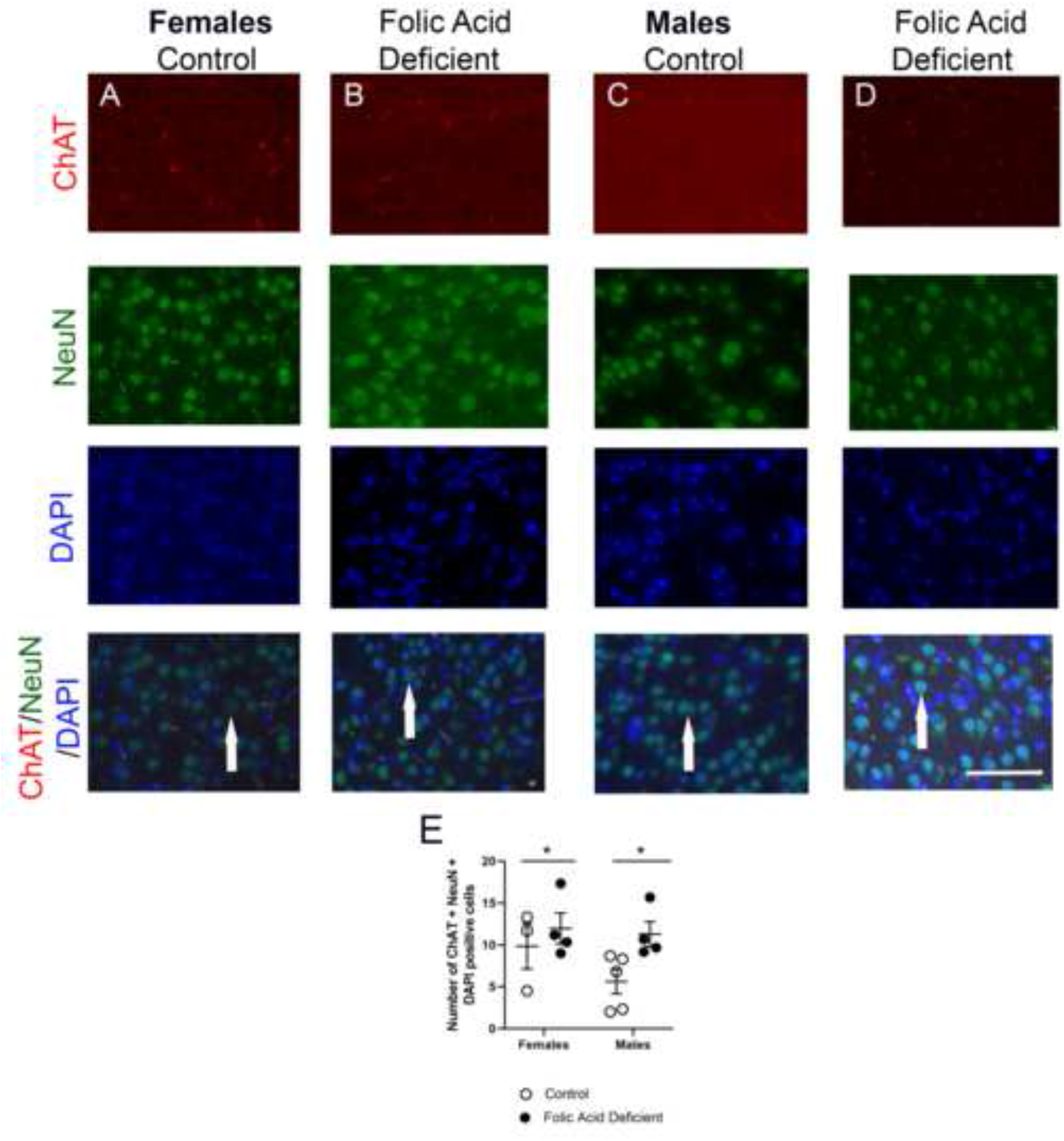
The impact of dietary folic acid deficiency and sex on neuronal choline acetyltransferase (ChAT) levels within the cortex of adult mice. Representative images from all experimental groups (A to D) of ChAT, neuronal nuclei (NeuN), and 4’,6-diamidino-2-phenylindole (DAPI). Quantification of ChAT, NeuN, and DAPI positive cells (E). Scatter plot with mean + SEM of 3 to 5 mice per group. * p<0.05 indicate Tukey’s pairwise comparison between groups.

To confirm changes in choline metabolism in brain tissue, we also measured acetylcholine esterase (AchE) levels in neurons within the cortex. Representative images are shown in Figure 5A to D. Quantification revealed no difference between sexes (F (_1,12_) = 0.49, p = 0.49), but did reveal decreases in FD mice compared to CD animals (Figure 5E; F(_1,12_) = 7.29, p = 0.019).

**Figure 5.**
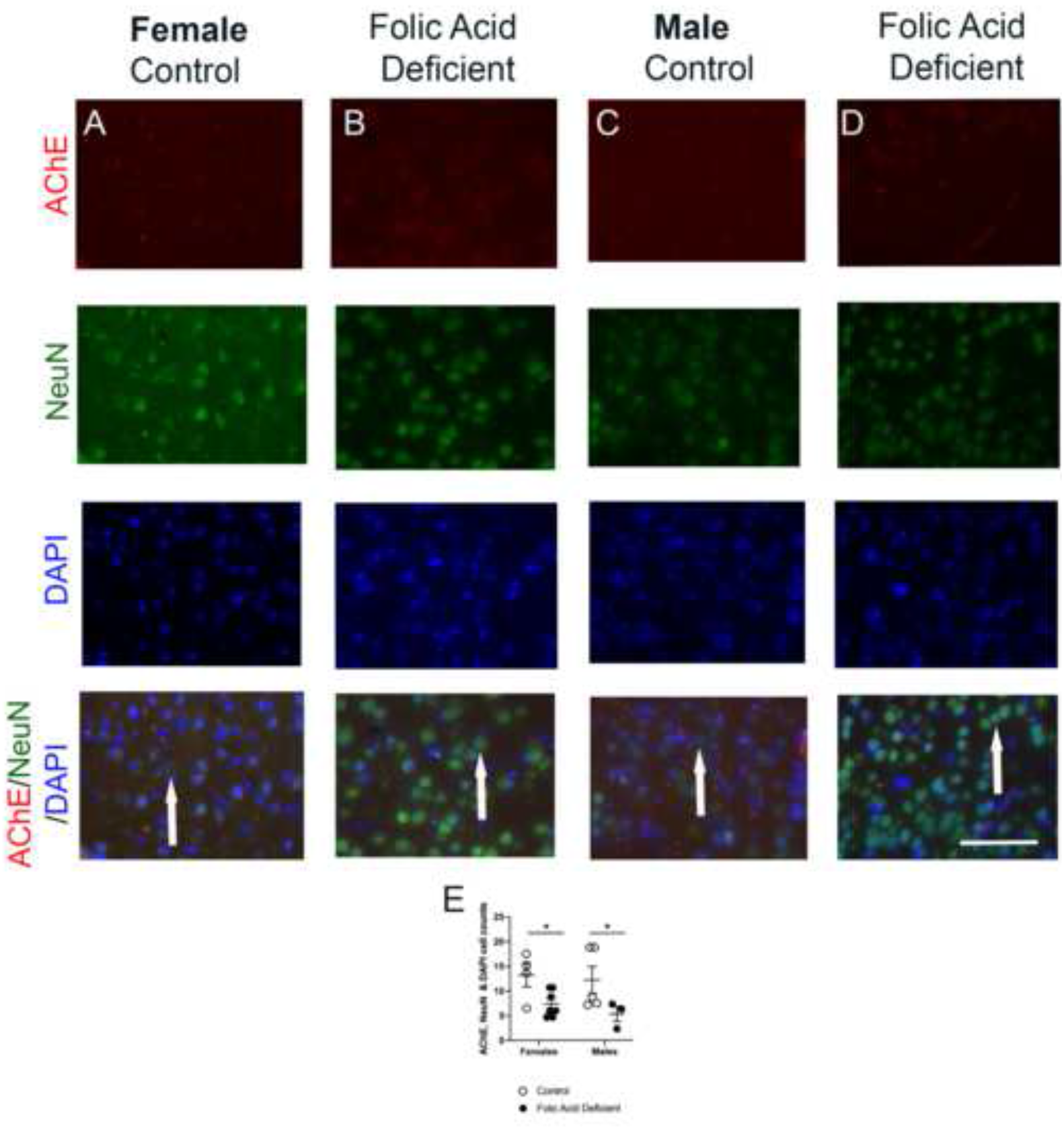
The effect of dietary folic acid deficiency and sex on neuronal acetylcholine transferase (AChE) levels within the cortex of adult mice. Representative images from all experimental groups (A to D) of AChE, neuronal nuclei (NeuN), and 4’,6-diamidino-2-phenylindole (DAPI). Quantification of AChE, NeuN, and DAPI positive cells (E). Scatter plot with mean + SEM of 3 to 5 mice per group. * p<0.05 indicate Tukey’s pairwise comparison between indicated groups.

## 4. Discussion

One-carbon metabolism involves multiple complex factors that coordinate gene expression and metabolism in response to nutritional signals that impact cellular anabolism, redox homeostasis, and epigenetics. There is growing evidence that sex plays a significant role in one-carbon metabolism and in regulating levels of blood homocysteine (22–25). Despite this, there is limited evidence for sex differences in brain tissue. Using a mouse model, our study investigated the impact of sex and dietary folic acid deficiency on hippocampal morphology and brain choline metabolism. As expected, the FD mice had lower serum folates than CD mice. Notably, our results indicate a significant decrease in the thickness of the granular layer within the CA1-CA3 region in both males and females maintained on a FD. In cortical tissue we demonstrate changes in acetylcholine metabolism between diet groups. The only sex difference we report in our study is the decreased apoptosis observed in the CA3 region of the hippocampus in males compared to females.

Consistent with prior research, our results demonstrate an intrinsic link between choline and folate metabolism (14–17). Specifically, our results show the novel finding of increased levels of cortical neuronal ChAT, the enzyme that catalyzes the synthesis of acetylcholine from choline at the synapse and decreased levels of AChE, the enzyme that catalyzes the destruction of acetylcholine within cortical neurons. Choline can act as a 1C donor in the liver and kidney via betaine homocysteine methyltransferase, especially when folate levels are low (33,34). This compensatory function of choline shunts choline away from the production of acetylcholine, potentially leading to lower levels of the neurotransmitter (14,15,17). Therefore, we suggest a potential mechanism for the demonstrated changes in cortical choline metabolism is a compensatory upregulation of ChAT and downregulation of AChE in response to decreased levels of acetylcholine. Low levels of acetylcholine have been associated with cognitive impairments, including mild cognitive impairment and dementia (35). An interesting follow-up to this study would be to assess cognitive function in adult females and males maintained on folic acid deficient and control diets.

The hippocampus is impacted by a genetic or dietary deficiency in 1C metabolism, as shown by others; our results demonstrate this effect as well (14,16,21). The different areas of the hippocampus and their functions are topics of ongoing research. In humans, the CA3 region has been specifically linked to spatial representations and episodic memory. Additionally, the CA3 region is susceptible to neurodegeneration, possibly accounting for our results (36). Both *in vivo* and *in vitro* studies have demonstrated that reduced levels of folate result in an increase in apoptotic markers (19,37–39), possibly due to increases in oxidative stress (40), DNA damage (38), or neuroexcitatory mechanisms (41). Recent *in vivo* studies have indicated similar results, with hyperhomocysteinemia, a direct consequence of 1C deficiency, leading to increased autophagy and apoptosis of cortical neurons(42).

While our results did not indicate a diet difference in apoptosis, there was a significant sex difference in apoptosis in the CA3 region of the hippocampus. The CA3 region is the most interconnected region of the hippocampus and has been implicated in memory functions and neurodegeneration (36). In order to measure levels of apoptosis, we stained for active caspase-3, which, as an executioner caspase leading to cell death, acts as an indirect measure of apoptosis (43). Previous research demonstrates a distinct sex difference in the activation of caspases, specifically following brain injury (44,45). These differences could be accounted for by sex hormones, potentially via the pro-apoptotic protein Bax (46). Model studies have indicated a significant sex difference in gross hippocampal morphology within the hippocampus including the dentate gyrus, CA1 and CA3 regions (47). A recent human study indicates that after adjusting for total hippocampal volume, regional sex differences exist, however, they are not present in the CA2/CA3 regions or the dentate gyrus (48). We are not observing a diet impact on apoptosis, which has been previously reported (19). We think that the duration and time point of dietary deficiency may be an important factor, since there is minimal apoptosis that occurs in the normal adult brain (49). Apoptosis in the brain is associated with key time points, such as after birth (49) and damage (50,51).

In conclusion, our study demonstrates stronger evidence that folate availability modulates changes in brain tissue more than sex in adult mice. Future research should aim to include behavioral analysis measuring cognitive function of male and females maintained on a 1C deficient diets.

## Acknowledgments

The authors would like to the College of Graduate Studies for the generous support of this collaboration. CB was funded by the Kenneth A. Suarez Research Fellowship.

## Notes on contribution

Calli Bennett BSc is a Doctor of Osteopathic Medicine student at Midwestern University. Calli graduated with an Honors Bachelor of Science degree in Anthropology from the University of Utah.

Jacalyn Green PhD is a Professor in the Department of Biochemistry and Molecular Genetics at Midwestern University Downers Grove campus.

Mae Ciancio PhD is an Associate Professor in the Biomedical Sciences Program at Midwestern University Downers Grove campus.

Joanna Goral PhD is an Associate Professor in the Department of Anatomy at Midwestern University Downers Grove campus.

Lenore Pitstick BS is a Department Research Coordinator in the Department of Biochemistry and Molecular Genetics at Midwestern University Downers Grove campus.

Matt Pytynia BA is a Senior Research Specialist in the Biomedical Sciences Program at Midwestern University Downers Grove campus.

Alice Meyer MBS is a Senior Research Associate in the Department of Anatomy at Midwestern University Downers Grove campus.

Neha Kwatra BSc is a Doctor of Dental Medicine student at Midwestern University. Neha graduated with a BSc in Physiology from the University of Arizona.

Nafisa M. Jadavji PhD is an Assistant Professor in Biomedical Sciences at Midwestern University (US) and Research Assistant Professor in Neuroscience at Carleton University (Canada).

